# Circadian photoperiod alters TREK-1 channel function and expression in dorsal raphe serotonergic neurons

**DOI:** 10.1101/2020.06.24.169532

**Authors:** Manuel A. Giannoni-Guzmán, Anna Kamitakahara, Valerie Magalong, Pat Levitt, Douglas G. McMahon

**Affiliations:** Department of Biological Sciences, Vanderbilt University, Nashville, TN, USA; Department of Pediatrics and Program in Developmental Neuroscience and Neurogenetics, The Saban Research Institute, Children’s Hospital Los Angeles, Keck School of Medicine, University of Southern California, Los Angeles, CA 90027, USA; Vanderbilt Brain Institute, Vanderbilt University, Nashville, TN, USA

## Abstract

Seasonal daylength has been linked to the development and prevalence of mood disorders, however, the neural mechanisms underlying this relationship remain unknown. Previous work in our laboratory has shown that developmental exposure to seasonal photoperiods has enduring effects on the activity of mouse dorsal raphe serotonergic neurons, their intrinsic electrical properties, as well as on depression and anxiety-related behaviors. Here we focus on the possible ionic mechanisms that underlie the observed photoperiodic programming of the electrophysiological properties of serotonin neurons, focusing on the twin-pore K+ channels TREK-1 and TASK-1 that set resting membrane potential and regulate excitability. Using multielectrode array recordings in *ex vivo* dorsal raphe slices, we examined the effects of pharmacological inhibition of these channels on the spike rates of serotonin neurons of mice from different photoperiods. Pharmacological inhibition of TREK-1 significantly increased spike frequency in Short and Equinox photoperiod cohorts, but did not further elevate the firing rate in slices from Long photoperiod mice, suggesting that TREK-1 function is reduced in Long photoperiods. In contrast, inhibition of TASK-1 resulted in increases in firing rates across all photoperiods, suggesting that it contributes to setting excitability, but is not regulated by photoperiod. To examine if photoperiod impacts transcriptional regulation of TREK-1, we quantified *Kcnk2* mRNA levels specifically in dorsal raphe 5-HT neurons using triple-label RNAscope. We found that Long photoperiod significantly reduced levels of *Kcnk2* in serotonin neurons co-expressing *Tph2*, and *Pet-1*, Photoperiodic effects on the function and expression of TREK-1 were blocked in melatonin 1 receptor knockout (MT-1KO) mice, consistent with previous findings that MT-1 signaling is necessary for photoperiodic programming of dorsal raphe 5-HT neurons. Taken together these results indicate that photoperiodic regulation of TREK-1 expression and function plays a key role in photoperiodic programming the excitability of dorsal raphe 5-HT neurons.

## Introduction

External environment signals can have significant impact on the development of brain and behavior. Early-life experiences have enduring effects on neural development, which in turn impact behavioral phenotypes in adulthood^1–4^. Day length (photoperiod) is a pervasive environmental signal that has been shown to program function of serotonergic function and behaviors in rodents^1,5,6^, and recently has been shown to associate with the life-time risk of major depression in humans^7^. Nevertheless, the neural and molecular correlates underlying this complex relationship remain to be found.

Perinatal photoperiod has been linked to enduring changes in the circadian clock expression in the suprachiasmatic nucleus (SCN), the central pacemaker of circadian activity^8^. More recently, our laboratory has shown that mice raised until weaning in long (summer-like) photoperiods exhibited less depressive- and anxiety-like behaviors as adults than mice raised in short (winter-like) conditions^5^. At the neural level, 5-HT neurons from Long mice showed increased excitability - depolarization of the resting membrane potential and a decrease in the amplitude of the afterhyperpolarization (AHP), both of which may contribute the increased spike frequency generated by a given amount of injected depolarizing current. Furthermore, these behavioral and neural changes elicited by developmental photoperiod were negated in melatonin receptor 1 (MT-1) knockout mice, indicating that melatonin signaling is required for the changes induced by photoperiod to DRN serotonin neurons. Although it is clear that developmental photoperiod programs 5-HT neurons, the specific molecules and processes that are targeted remain to be elucidated.

Resting membrane potential in many neurons is maintained by twin-pore K+ channels. These channels shape the excitability of neurons by setting the resting membrane potential and by opposing depolarizing inputs^9^. In the mouse DRNs the most highly expressed isoforms of these channels are TREK-1 and TASK-1^10^. Given the changes induces by developmental photoperiod both at the behavioral and neural levels, examining the response of these channels in mice to photoperiod may provide insight into the mechanisms of photoperiodic programming of 5-HT DRN neurons.

TREK-1 is of particular interest since the removal of the channel via constitutive knockout results in an anti-depressive-like phenotype^11^. TREK-1 is known to be regulated by the induction of cAMP fluctuations thru Gs- or Gi-coupled receptors and activation of Gq protein pathways^12,13^. *In vivo* extracellular recordings of DRN 5-HT neurons revealed that a significant increase in the firing rate of neurons of TREK-1 KO compared to wild type mice^11^. These findings suggest TREK-1 channel as a novel candidate target for the treatment of depression^14^.

TREK-1 channels are mostly insensitive to antagonists of voltage-gated K+ channels such as tetraethylammonium (TEA) and 4-aminopyridine (4-AP)^15^, however, recent work has shown the successful inhibition of TREK-1 using amlodipine, a dihydropyridine originally used Ca2+ channel antagonist^16^. The IC50 of amlodipine for TREK-1 inhibition (430nM) is 100-fold less than the required dose for inhibition of Ca2+ channels, making one of the first specific antagonists to the channel^17–19^. More recent work has shown that spadin, a peptide derived from the maturation of the neurotensin receptor 3, specifically binds to TREK-1, blocking channel activity *in vitro* and increases the firing rate of 5-HT neurons of the raphe *in vivo*^20^. Also, in behavioral tests spadin-treated mice presented depression resistant phenotypes, similar to TREK-1 KO.

The primary goal of this study was to determine the potential roles of TREK-1 and TASK-1 in the ionic mechanisms that alter the spontaneous spike rate of dorsal raphe 5-HT neurons in mice raised in different photoperiods. We hypothesized that photoperiod increased excitability of 5-HT neurons could result from a reduction in TREK-1 and/or TASK-1 activity. Furthermore, potential changes to the response of pharmacological inhibition could be the result of changes in the gene expression profile of the channel. To test these hypotheses, we performed multi-electrode array recordings on Long, Equinox and Short photoperiod mice were we pharmacologically inhibited either TREK-1 and TASK-1 channels. In addition, we examined the mRNA expression levels of *knck-2* (TREK-1) in 5-HT neurons using quantitative *in situ* hybridization technique RNAscope.

## Results

### Photoperiod alters DRN 5-HT Neuron Spike Rate through TREK-1 but not TASK-1

To determine if photoperiod programs TREK-1 or TASK-1 in DRN serotonin neurons we performed multielectrode array recordings on acute brain slices containing the dorsomedial region of the DRN of mice (C3Hf+/+) conceived and raised in either Long (summer-like), Equinox (spring/fall like) or Short (winter-like) photoperiods, and tested for dose-response effects of specific pharmacological inhibitors on spike rate. Serotonin neurons were identified by suppression of spiking in response to 8-OHDPAT at the end of each recording. Data were analyzed by Repeated Measures Two-Way ANOVA for main effects on spike rate of photoperiod, inhibitor dose, and an interaction of photoperiod with inhibitor dose. We hypothesized that if one of these K+ channels contributes to by photoperiodic programming of excitability, then this would be revealed by a significant interaction between photoperiod and inhibitor dose on spike rate.

TREK-1 channels were pharmacologically inhibited using the dihydropyridine amlodipine besylate (0-10uM, IC50 = 430nM) which has been shown to be a potent inhibitor of TREK-1 channels at concentrations where it has minimal antagonism of Ca2+ channels^16^. We hypothesized that photoperiod increased excitability of 5-HT neurons might result from a reduction in TREK-1 activity, thus 5-HT neurons from Long photoperiod mice would be unresponsive to pharmacological inhibition of TREK-1. Consistent with this hypothesis, the firing rate of 5-HT neurons from the Long cohort was not increased by amlodipine besylate, even at doses well-above the IC50, while that of neurons originating from Equinox and Short photoperiod mice increased in a dose dependent manner (Figure 1A). Repeated Measures Two-way ANOVA analysis revealed a significant interaction between photoperiod and amlodipine dose (*p* < 0.001***) and main effects of amlodipine dose alone (*p* < 0.0001****). At the higher doses amlodipine may have partially inhibited Ca2+ currents, so we used a second specific TREK-1 inhibitor spadin (0-7uM, IC50 = 70.7nM) to further test our hypothesis. Results with spadin were similar to amlodipine, with no further effect of spadin inhibition of TREK-1 on the already elevated spike rate of neurons of Long photoperiod mice (Figure 1B). Overall the RM Two-Way ANOVA analysis revealed a significant interaction between photoperiod and spadin dose (*p*<0.001***). As with amlodipine we also observed a significant increase in the spike rate of the Equinox cohort, but the increase in firing rate in the Short cohort with spadin did not reach statistical significance. These results suggest that photoperiod programs TREK-1 channel function in dorsal raphe 5-HT neurons.

**Figure 1.**
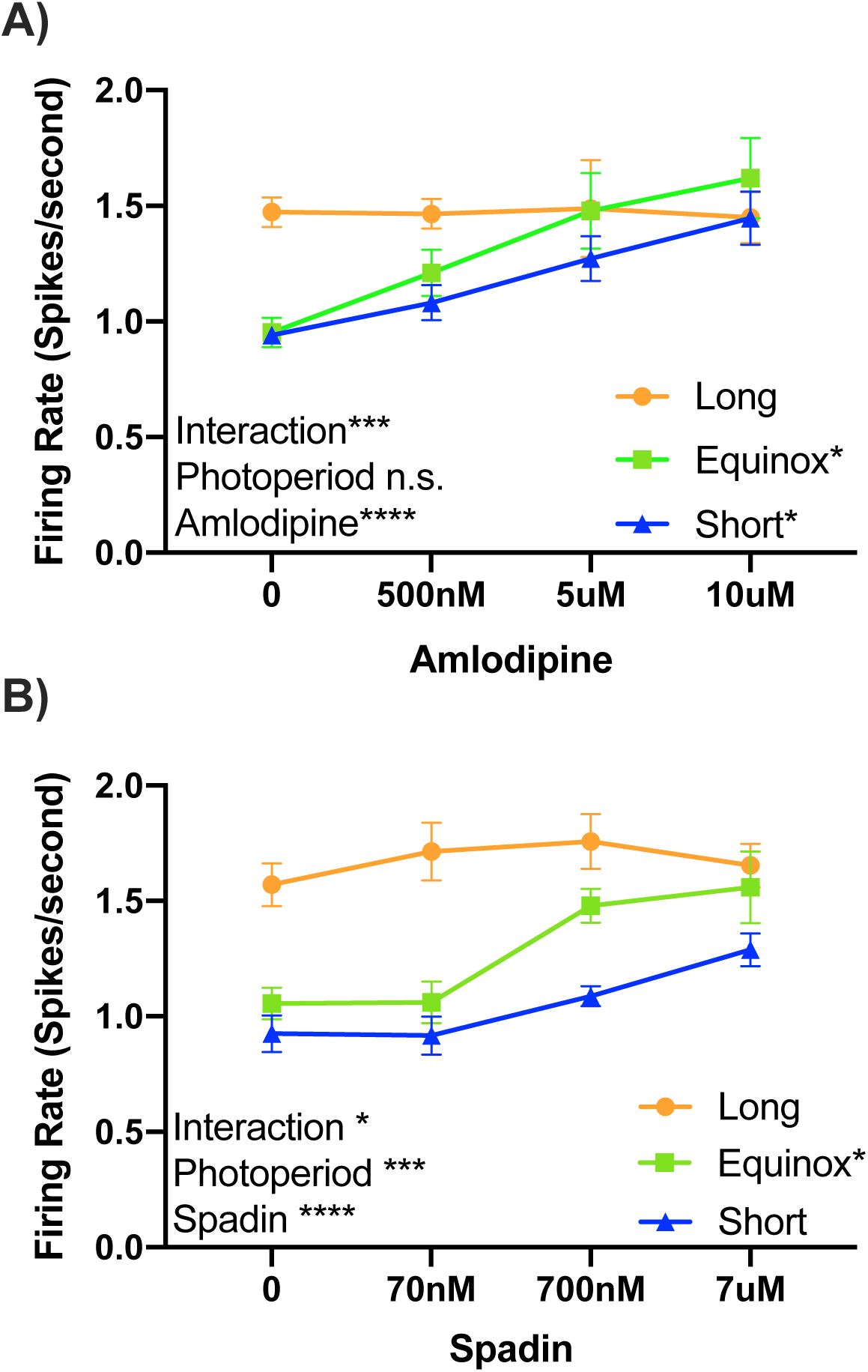
TREK-1 function is modulated by photoperiod. **A)** Firing rate of dorsal raphe 5-HT neurons exposed to increasing doses of amlodipine from Long, Equinox and Short photoperiod mice. Two-Way analysis of variance (ANOVA), showed a significant effect for dose of Amlodipine (*p*<0.0001****) and the interaction of Amlodipine and Photoperiod (*p*=0.0008***). Tukey’s multiple comparisons revealed a significant increase in firing rate in response to amlodipine in the Equinox (*p*=0.0184*, 0 vs. 500nM) and Short groups *(p=*0.0366*, 0 vs. 10uM). **B)** Pharmacological inhibition of TREK-1 with spadin revealed significant main effects in dose of spadin (*p*=0.0002****) and Photoperiod (*p*<0.0001***), as well as a significant interaction (*p*=0.0188*). Tukey’s multiple comparisons revealed a significant increase in firing rate only in the Equinox group (*p*=0.0147*, 0 vs. 700nM).

In addition to TREK-1, the TASK-1 channel is also highly expressed in the DRN and shares the role of maintaining the resting membrane potential. Using a highly specific TASK-1 inhibitor ML-365 (0-10uM, IC50 = 4nM,^21^), we tested if photoperiodic programming altered TASK-1 function in serotonin neurons of the DRN. TASK-1 inhibition resulted in a significant increase in firing rate in the three photoperiod cohorts (Figure 2). Statistical analysis via two-way RM ANOVA revealed significant effects of Ml-365 dose (*p*<0.0001****) and photoperiod (*p* = 0.004**), but no significant interaction (*p* = 0.68). These data suggest that while TASK-1 can regulate the excitability in DRN serotonin neurons, it does not contribute to photoperiodic programming.

**Figure 2.**
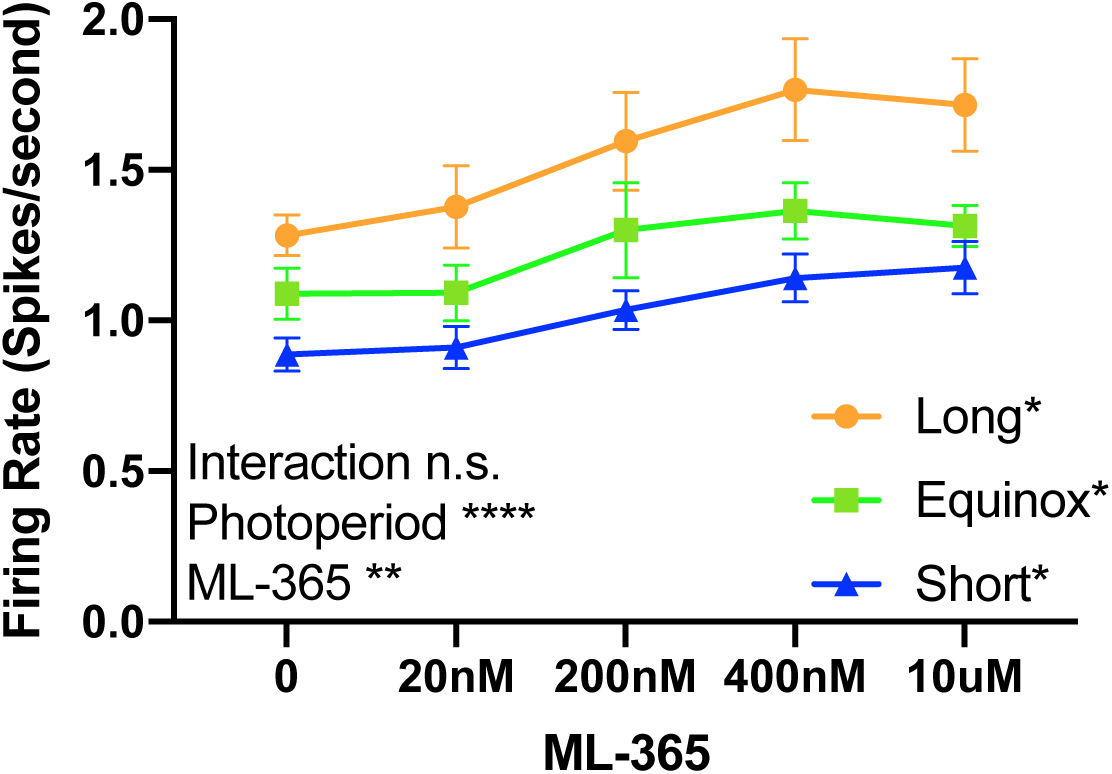
TASK-1 function regulates spike rate, but is not modulated by photoperiod. **A)** Firing rate of dorsal raphe 5-HT neurons exposed to increasing doses of selective TASK-1 inhibitor ML-365 from Long, Equinox and Short photoperiod cohorts. Two-way ANOVA showed a significant effect for dose of ML-365 (*p*<0.0001****), Photoperiod (*p*=0.0042**), while there was no significant interaction (*p*=0.6818). Tukey’s multiple comparisons revealed a significant increase in firing rate in response to ML-365 in all groups (Long: *p*=0.0328* 0 vs. 20nM; Equinox *p=*0.0497*, 0 vs. 400nM; Short: *p*=0.0392, 0 vs. 400nM).

### Long Photoperiod Decreases the Expression TREK-1 mRNA in Serotonin Neurons

To determine if the decrease in measurable TREK-1 inhibition in Long photoperiod was the result of changes at the gene expression level, we employed RNAscope to determine the abundance of *Kcnk2* exclusively in *Tph2* and *Pet-1* positive neurons of the DRN (Figure 3). The resulting analysis revealed significant differences in the levels of *Kcnk2* (TREK-1) transcripts colocalized with *Tph2* and *Pet-1* (One Way ANOVA, F = 4.991, *p* = 0.0218). Specifically, 5-HT neurons of Long photoperiod mice had significantly lower levels of *Kcnk2* transcripts, compared to Short photoperiod mice (Figure 3).

**Figure 3.**
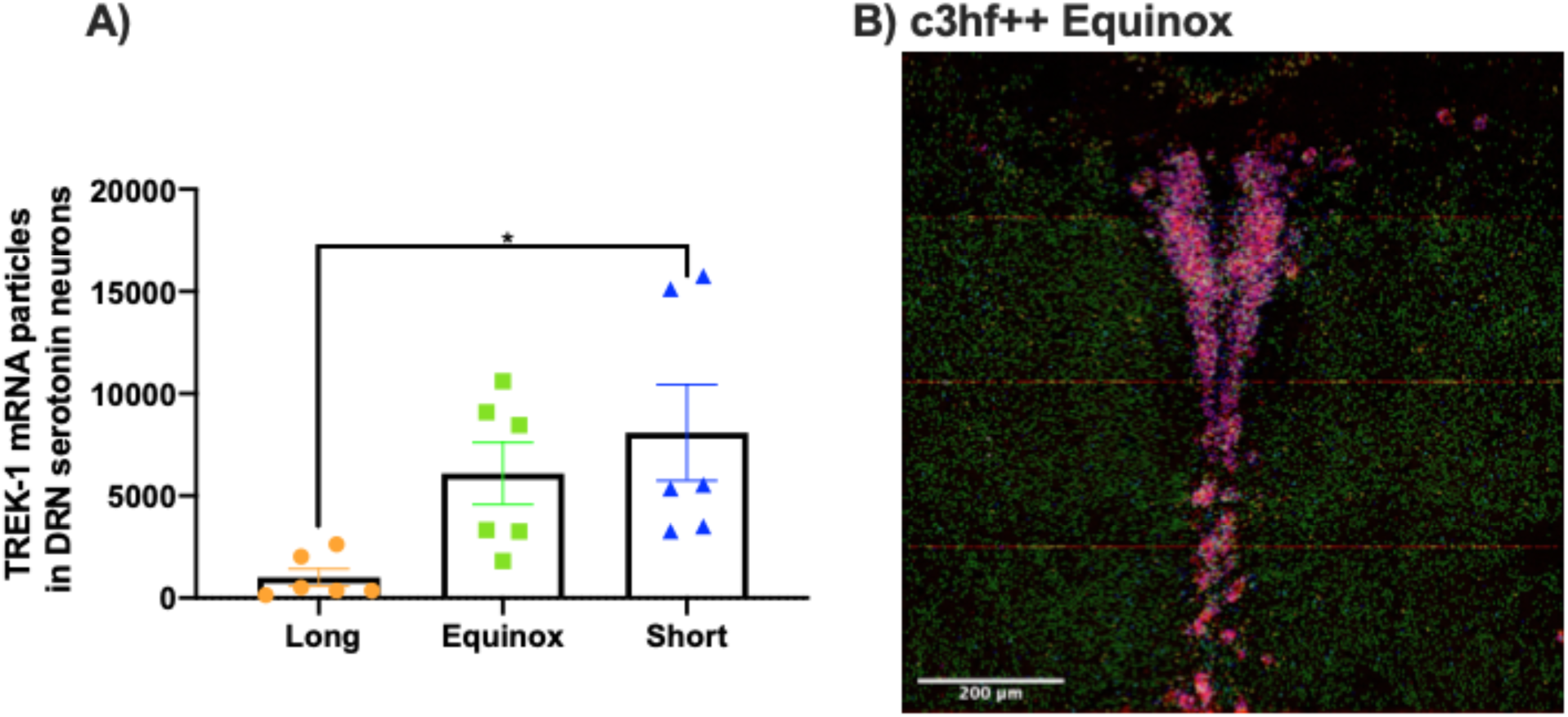
mRNA levels of *Kcnk2* (TREK-1) in *Tph2* and *Pet-1* positive neurons are decreased in Long photoperiod mice. Quantification of colocalized *Kcnk2, Tph2* and *Pet-1* in the dorsal raphe. **A)** Colocalized transcript detection of *Kcnk2* in c3hf++ mice raised in long equinox or short photoperiods revealed a significant decrease of the transcript in serotonin neurons of long photoperiod mice (One Way ANOVA, F=4.991, *p*=0.0218*). Tukey’s post hoc multiple comparison revealed significant differences between Long and Short photoperiod cohorts. **B)** Representative image of c3hf++ dorsal raphe from a mouse from the Equinox photoperiod, showing quantification of colocalized transcripts. The image was made from 16 captures at 40x. *Kcnk2* (green), *Tph2* (red) and *Pet-1* (blue), colocalization is shown of *Pet-1* and *Tph2* is in purple, *Kcnk2* colocalization is in either turquoise or yellow.

### Photoperiod programming of TREK-1 requires melatonin signaling

Photoperiodic programming of the excitability of DRN serotonin neurons requires melatonin signaling through the MT-1 melatonin receptor^5^. Given our results suggesting that TREK-1 is a target of this programming, we hypothesized that in the absence of a functional melatonin pathway, TREK-1 would not be regulated across photoperiods. We tested this using spike rate assays of amlodipine inhibition of TREK-1 in DRN slices, and RNAscope of TREK-1 expression in DRN serotonin neurons. In contrast to the above results for mice with intact melatonin signaling (Figure 1), in MT-1 knockout mice TREK-1 inhibition by amlodipine had no significant effect on the firing rate of DRN serotonin neurons in any photoperiod, and there was no interaction between photoperiod and amlodipine (Figure 4A). Furthermore, in MT-1 knockout mice DRN serotonin neuron *Kcnk2* transcript expression was similar across all photoperiods (Figure 4B), again in contrast to the observed decrease in *Kcnk2* in Long photoperiod melatonin signaling-competent mice (Figure 3). These results suggest that photoperiod acts on TREK-1 expression and function through MT-1 melatonin signaling to program DRN serotonin neuron excitability.

**Figure 4.**
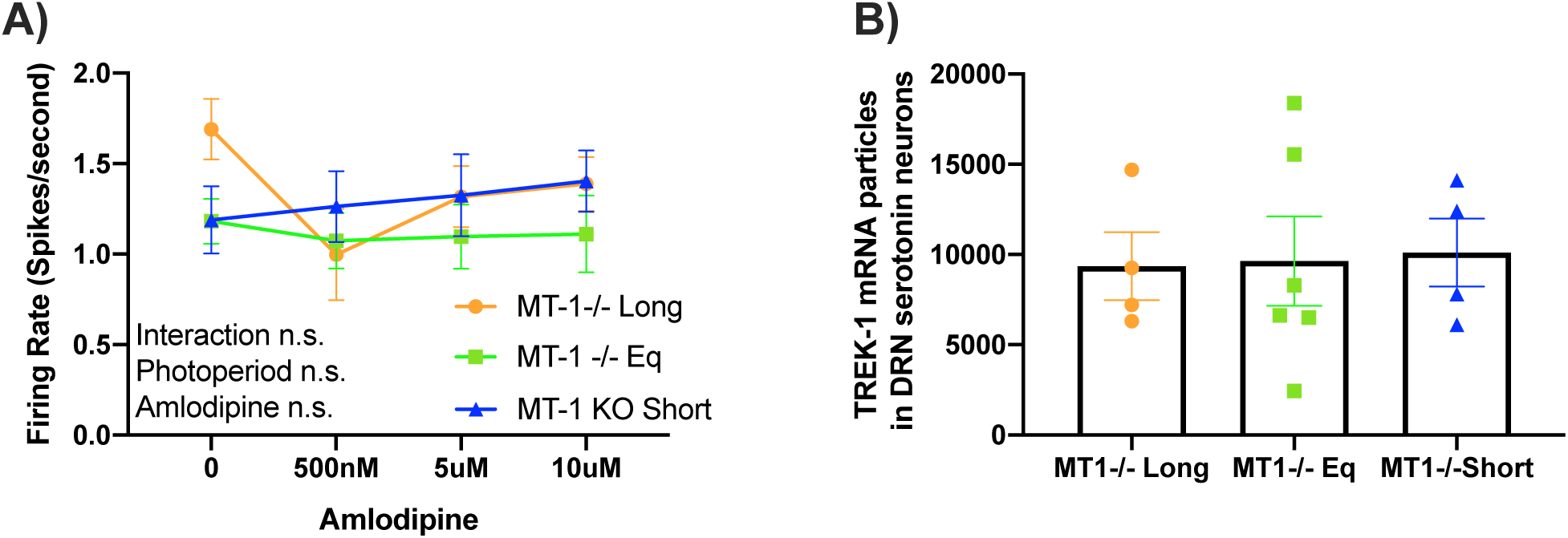
Melatonin receptor 1 knockout abolishes photoperiodic programming of TREK-1 in serotonin neurons. **A)** Firing rate of dorsal raphe 5-HT neurons exposed to increasing doses of Amlodipine from MT-1 KO mice raised in Long, Short or Equinox photoperiods. Two-Way analysis of variance (ANOVA), did not reveal significant changes between the cohorts. **B)** Mean *Kcnk2* expression levels in 5-HT neurons from MT-1 KO mice were similar irrespective of photoperiod.

## Discussion

Our findings demonstrate that photoperiod differently alters the function and expression of the key resting membrane potential regulator TREK-1 in adult 5-HT neurons, but not TASK-1. Results with TREK-1 blockers amlodipine and spadin, suggest that reduced TREK-1 function following exposure to Long photoperiods is a key element in the photoperiod-driven increase in 5-HT neuron spike rate and excitability, and quantification of *Kcnk2* transcript levels by RNAscope demonstrated directly that TREK-1 expression is reduced in DRN serotonergic neurons in Long photoperiods. Finally, we tested the necessity of melatonin MT-1 signaling for photoperiod programming of TREK-1, and found that, like programming of overall DRN serotonin neuron excitability, the programming of TREK-1 function and expression is dependent on melatonin signaling. Taken together, our findings suggest that photoperiod acts at the transcriptional level on TREK-1, potentially influencing the number of active TREK-1 channels on 5-HT neurons, which then contributes to the previously observed differences in excitability of these neurons as well as anxiety-like and depressive-like phenotypes following development in different photoperiods.

Research examining TREK-1’s association with depression-like behaviors has shown that deletion of the sorting protein NTSR3/Sortilin (Sort1-/-) results in depression-resistant phenotypes, increased activity of DRN neurons, and a decrease in cell surface levels of TREK-1. Sort1-/- mice present similar phenotypes and electrophysiological properties of 5-HT neurons to TREK-1 knockout mice and Long photoperiod mice^5,11,22^. Thus, the lack of significant increase in firing rate in Long photoperiod mice upon TREK-1 inhibition suggests low levels of functional TREK-1 channels in 5-HT neurons. Furthermore, spadin, which we used to inhibit TREK-1, when administrated to mice via intraperitoneal injections elicits an increase in the firing rate of 5-HT neurons *in vivo* and a decrease in depression-like behaviors^20,23^. Thus, our results provide strong support for programming of TREK-1 channels as a key driver of the behavioral effects of developmental photoperiod.

Our finding that *Kcnk2* levels in 5-HT neurons are significantly decreased in Long photoperiod mice provides a potential mechanism by which photoperiod modulates TREK-1 channels, but it does not preclude other potential avenues. For example, it is possible that photoperiod may also be acting on NTSR3/Sortilin, which would affect the trafficking of TREK-1 to the membrane. This is a possible scenario in the context of our Short photoperiod results where we observed small increases in firing rate in response to spadin, but *Kcnk2* transcript levels are high in 5-HT neurons (Figures 1,3).

Consistent with previous results showing that melatonin signaling is required for photoperiodic programming of 5-HT neurons in the raphe^5^, our results here show that MT-1 knockout mice are irresponsive to photoperiod, and that MT-1 knockouts are unresponsive to pharmacological inhibition of TREK-1 (Figure 4). Thus, it is possible that the lack of melatonin signaling in these mice keeps the 5-HT system from adjusting to photoperiod. In C57BL/6 mice TREK-1 is expressed in different regions of the brain as early as E15, and expression levels assayed thru in situ hybridization and qPCR, remain stable thru adulthood^24^. Since C57BL/6 mice are not melatonin-competent, it would be interesting to examine the levels of TREK-1 expression across different photoperiods in melatonin competent mice (e.g. C3Hf+/+) throughout embryonic development.

The work presented here shows that an external environmental cue, daylength, programs a critical component of the neuronal machinery of serotonin neurons at the transcriptional level, thus causing changes of the electrophysiological properties of these neurons, which can lead to enduring changes in depression and anxiety behaviors. Understanding the mechanisms of TREK-1 channel modulation in relation to serotonergic function and behavior may lead to new treatments for mood disorders. Our future efforts will concentrate on dissecting how developmental photoperiod via melatonin signaling pathways regulates TREK-1.

## Methods

### Animals and Housing

C3Hf^+/+^ males and females were developed and raised to maturity on either Equinox (12 hours of light 12 hours of darkness), Short (8 hours of light and 16 hours of darkness) or Long (16 hours of light and 8 hours of darkness) photoperiods. Experimental assays were performed between P40 to P60 with tissues isolated at 11:00am-12:00pm local time, the mid-day point on all light cycles. Male and female animals were used in equal numbers in each cohort. MT-1 receptor knockout mice in the C3Hf+/+ background were bred and raised under the same conditions as the C3Hf^+/+^ mice. Genotype was confirmed before breeding the animals and before each experiment. Experiments were performed in accordance with the Vanderbilt University Institutional Animal Care and Use Committee and National Institutes of Health guidelines.

### Multielectrode Array Electrophysiological Recording

Mice were euthanized, brains were extracted and mounted in cold, oxygenated (95%O_2_-5%CO_2_) dissecting media (in mM: 114.5 NaCl, 3.5 KCL, 1 NaH2PO4, 1.3 MgSO4, 2.5 CaCl, 10 D(+)-glucose, and 35.7 NaCHO3), and 300μm thick coronal slices were taken using a Leica V2100 (Leica Instruments). Dorsal raphe nuclei were isolated by removing the extraneous cortical tissue and placed sample in a slice chamber full of room temperature, oxygenated, extracellular recording media (in mM: 124 NaCl, 3.5 KCl, 1 NaH2PO4, 1.3MgSO4, 2.5 CaCl2,10 D(+)-glucose, and 20 NaHCO3). Multielectrode array recordings were performed by dissecting out the dorsal raphe nucleus after slicing. The tissue was placed on a perforated electrode array and immobilized with a harp for recording. To elicit spontaneous firing of serotonin neurons 40 µM tryptophan and 3 µM phenylephrine were added to the recording solution which was perfused (1.3 ml/sec) over the slice. After 45 minutes of placing the slice in solution recording began. For TREK-1 inhibition amlodipine was perfused (1.3 ml/sec) over the slice in a serial manner at concentrations of 0, 500nM, 5µM and 10µM. Each dose was perfused for 5-min to ensure the desired concentration in the bath. For spadin the same protocol was used with the following dosages 0, 70nM, 700nM and 7µM. Inhibition of TASK-1 was achieved using highly specific inhibitor ML-365 at 0, 20nM, 200nM and 400nM^21^. Lastly, 8-OH DPAT was perfused into the solution at 5uM concentration to silence serotonergic cells.

## RNAscope quantification

Mouse (n = 6 per group) whole brain tissue was collected and submerged into isopentane for 25 seconds, placed in crushed dry ice for approximately 1 minute, and stored in a 50mL falcon tube at −80°C. 20 µm sections were prepared per animal targeting the dorsal raphe (DRN) from - 5.5mm to −5.75mm from Bregma. Tissue sections were then stained with RNAscope in situ hybridization, targeting the transcripts *Kcnk2* (TREK-1), *Tph2*, and *Pet-1* with fluorescent dyes Alexa488, Atto550, and Atto647, respectively. Once tissue was processed, laser scans for each fluorescent channel containing the dorsal raphe nucleus for each brain slice were captured using a Zeiss LSM510 confocal scanning microscope (Carl Zeiss Microscopy Gmbh, Jena, Germany). Sixteen sequential scans at 40x of 512×512 pixels were taken by the microscope and stitched together to produce a high-resolution composite of the area. Laser settings and pinhole sizes were the same for all the captures. Image analysis was performed using the open source software Fiji (https://imagej.net/Fiji). Colocalization of *Kcnk2* with *Tph2* and *Pet-1* was established using the plugin ComDet (https://github.com/ekatrukha/ComDet)^25^. Briefly, the plugin determines small ROIs each channel independently and determines if they colocalize spatially with up to 4 pixels of distance from each other. Number of colocalized particles is reported as a proxy for gene expression levels.

### Statistical Analysis

Traces were analyzed using offline sorting (Plexon) and spikes were sorted using a combination of manual identification and automatic K means based sorting software. Graphs and statistics were prepared using GraphPad PRISM 8. Repeated Measures Two-way ANOVA was performed for each of the cohorts using photoperiod and inhibitor dose as independent variables. Since we used male and female mice in each of the cohorts in equal numbers, sex differences were analyzed for all cohorts. No significant differences were observed in either C3Hf^+/+^ or MT-1KO cohorts.

## Acknowledgements

This research was supported by National Institutes of Health Grants NIH R01 MH108562 to DGM and 1P50MH096972 to the Vanderbilt University Conte Center. The authors would like to thank, Turnee Malik, Michael Tackenberg, David Sprinzen, and the Vanderbilt Vision Research Center for their advice and support.

